# Galvanic vs. Pulsatile Effects on Decision-Making Networks: Reshaping the Neural Activation Landscape

**DOI:** 10.1101/2023.09.01.555903

**Authors:** Paul Adkisson, Cynthia R. Steinhardt, Gene Y. Fridman

**Affiliations:** Johns Hopkins University; Columbia University

## Abstract

Primarily due to safety concerns, biphasic pulsatile stimulation is the present standard for electrical excitation of neural tissue with a diverse set of applications. While pulses have been shown to be effective to achieve functional outcomes, they have well-known deficits. Due to recent technical advances, galvanic stimulation, delivery of current for extended periods of time (>1s), has re-emerged as an alternative to pulsatile stimulation. In this paper, we use a winner-take-all decision-making cortical network model to investigate differences between pulsatile and galvanic stimulation in the context of a perceptual decision-making task. Based on previous work, we hypothesized that galvanic stimulation would produce more spatiotemporally distributed, network-sensitive neural responses, while pulsatile stimulation would produce highly synchronized activation of a limited group of neurons. Our results *in-silico* support these hypotheses for low-amplitude galvanic stimulation but deviate when galvanic amplitudes are large enough to directly activate or block nearby neurons. We conclude that with careful parametrization, galvanic stimulation could overcome some limitations of pulsatile stimulation to deliver more naturalistic firing patterns in the group of targeted neurons.

## II. INTRODUCTION

An accepted theoretical model of how we reach a decision given two alternatives posits that the average firing rates of two opposing neural populations encode the saliency of each alternative, and a decision is made when one of the two populations clears some firing rate threshold^1–3^. Using a well-accepted *in-silico* model of this process, we asked how electrical stimulation of one of the neural populations would influence the decision-making process. This problem is nontrivial because of the recurrent nature of the network with excitatory and inhibitory interconnections. We compared two forms of electrical stimulation each applied to a well-known decision-making task: conventional pulsatile stimulation and more recently re-introduced, galvanic stimulation.

Biphasic pulsatile stimulation (PS) is the standard for safe and effective electrical stimulation of the brain. In basic research, low-amplitude (e.g. 5µA) sub-millisecond pulses or sequences of pulses are commonly used to probe brain connectivity and function^4–7^. Clinically, pulses are used in neural prosthetics^8^, preresection surgeries for drug-resistant epilepsy^9^, deep brain stimulation for Parkinson’s disease^10^, and a host of emerging applications in “bioelectronic medicine”^11^. While pulses are clearly capable of inducing sensory percepts^8^, muscle movements^12^, and bias decision making^13^, there are clear limitations to their ability to deliver natural sensation^14^ or motion^15^. Evoked spikes are phase-locked to the pulse presentations^16^. The effect of pulse trains on an isolated neuron’s evoked spike rate varies in a nonlinear fashion. Pulses can facilitate and interfere with each other and spontaneous activity depending on pulse amplitude, pulse rate, and spontaneous baseline activity^17–19^. Therefore, neurons will respond differently depending on their distance from the electrode.

Galvanic stimulation (a.k.a. “direct current” or DC) (GS) has been limited to transcutaneous application (such as tDCS), due to safety implications associated with electrolysis and pH changes at the metal electrode^20,21^. This limitation however is being addressed in the recent development of implantable galvanic stimulation devices^21,22^ and conventional stimulators with high capacity electrodes^23^. Our understanding of how an implanted electrode delivering GS could affect a neural network is limited. GS has been shown to be capable of smoothly modulating a neuron’s firing rates up and down while preserving natural firing statistics and without producing unnatural synchrony due to phase-locking^24–27^. Biophysical modeling predicts that GS has these effects because it modulates axonal sensitivity to incoming synaptic inputs, smoothly up-modulating and down-modulating the firing rate from the spontaneous baseline^27–29^. Along with this smooth modulation, amplitude-dependent and polarity-dependent neural block makes the prediction of large network behavior more difficult^27,28,30^.

The single neuron effects of PS and GS lead to hypotheses of how a network of excitatory and inhibitory neurons would behave. Since GS modulates sensitivity to incoming synaptic activity, we hypothesize that a population of neurons experiencing GS will be more sensitive to network level excitation and inhibition than if this population were subjected to PS. The ability of a network to reach persistent activity states is critical in maintaining working memory for perceptual decision making^2^. We also hypothesize that GS activation will be spread more uniformly throughout the stimulated population, while PS may have high population-wide variability between high firing rate neurons and low firing rate neurons. Such variability in neural activation may contribute to the commonly observed variability across subjects in the effectiveness of cortical microstimulation^12–14^. We further hypothesize that PS will induce unnaturally high levels of neural synchrony which may lead to different decision-making latencies.

To explore these predictions, we extended a well-established computational model of perceptual decision making^2^ by adding pulsatile and galvanic stimulation to bias the decision-making output of the model. Both pulsatile and cathodic galvanic stimulation were capable of biasing behavior in favor of the stimulated population (excitatory bias). Anodic galvanic stimulation was also capable of biasing behavior against the stimulated population (inhibitory bias). In line with our hypothesis, the introduction of pulsatile stimulus caused the neurons in the vicinity of the stimulation electrode to respond with synchronous rapid firing, regardless of the state of the network, and cathodic galvanic stimulation caused more spatiotemporally distributed population responses with magnitudes that were highly dependent upon ongoing network activity. Contrary to our hypothesis, the increase in synchronous firing induced by PS did not lead to any differences in decision times relative to CGS.

## III. RESULTS

### Experimental Design

In this *in-silico* study, we assessed the ability of pulsatile (PS) and direct current (a.k.a. “galvanic”) stimulation (GS) to bias large networks of neurons by exposing a well-established computational model of perceptual decision making^2^ to both paradigms. In brief, this winner-take-all model consisted of two subpopulations of pyramidal neurons (P1 and P2) responsive to leftward and rightward dot motion respectively, non-selective pyramidal neurons (NS), and a common pool of inhibitory interneurons (Int), shown in Figure 1A. All neurons were simulated with leaky-integrate-and-fire (LIF) dynamics and connected with varying synaptic weights as in Wang^2^. We used the neural network parameters described in that publication to build our network. Neurons were given strong connections (*w_strong_*=1.7) within subpopulations (ex. P1 to P1), weak connections (*w_weak_*=0.8765) across subpopulations (ex. P1 to P2), and medium-strength connections (*w_medium_*=1) with interneurons. In each of 100 trials, all neurons received 2400Hz background Poisson-random excitatory input (AMPA EPSCs) resulting in spontaneous firing rates of 0-4spk/s for pyramidal neurons and 5-7spk/s for interneurons (Figure 1B). During the experiment, from t=1-3s, P1 and P2 received differential task input (AMPA EPSCs) varying from 0-80Hz depending on the task coherence for that trial (see Methods for details). For the example in Figure 1, when the task coherence is −51.2%, P2 neurons receive task input EPSCs at 60Hz and P1 neurons receive task input EPSCs at 20Hz. For this control experiment, this was the only input to the model. As a result of this input, P1 and P2 firing rates increased until P2 exceeded a natural threshold of about 15spk/s, and subsequently P2 firing rates rapidly increased, winning the decision. Subsequently, P1 firing rates were suppressed by feedforward inhibition from the interneurons (Figure 1B). Consistent with previous work, we defined “winning” and “losing” by comparing the average firing rates of the pyramidal populations P1 and P2 at the end of each trial (see Methods for details). If, during this period (t=1-3s) we exposed neurons in P1 to electrical stimulation, we could bias the network behavior, reversing the decision outcome, making P1 win and suppressing the P2 response (Figure 1C). After task input and electrical stimulation were turned off, the high-firing-rate attractor state was self-perpetuated by the network in both control and stimulation trials as seen between 3 and 4 seconds in Figures 1B and 1C, respectively.

**Figure 1.**
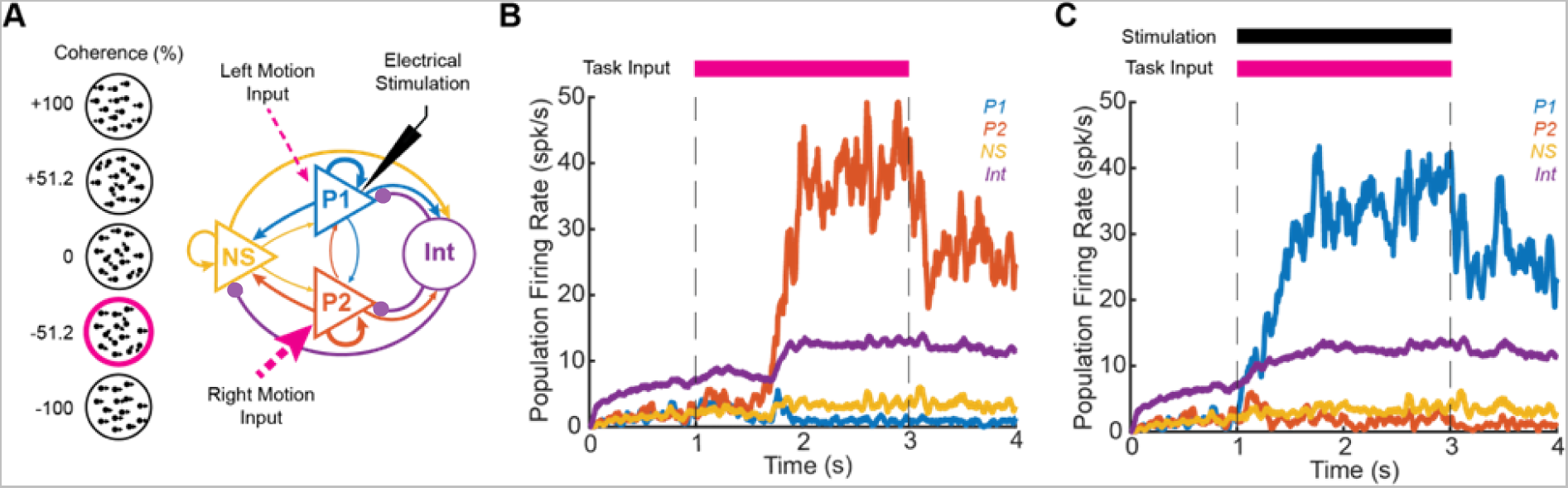
Experimental Design. (A) Model consists of two subpopulations (P1 and P2) responsive to leftward and rightward motion, non-selective pyramidal neurons (NS) and inhibitory interneurons (Int). Neurons are connected with strong, medium, and weak connections (thickness proportional to strength), with purple arrows indicating inhibitory connections. During a trial, all neurons receive background input. From 1-3s, P1 and P2 receive task-related input proportional to coherence of left versus rightward motion. P1 also receives electrical stimulation from 1-3s (black) to bias the network behavior. (B) Mean population firing rates of P1 (blue), P2 (red), NS (yellow), and Int (purple) in a representative control trial without electrical stimulation. Task input strongly favors P2 (−51.2% coherence) resulting in P2 winning the trial. (C) Mean population firing rates in a representative trial with pulsatile stimulation of P1. Despite task input favoring P2 (−51.2% coherence), pulsatile stimulation biases the network such that P1 wins the trial.

### Electrical Stimulation Alters Decision Making

The electrical stimulation paradigms were point-source monopolar (referenced to distant ground) pulsatile charge balanced biphasic pulse trains (PS), and cathodic (excitatory) and anodic (inhibitory) galvanic currents (CGS and AGS respectively). The stimulation paradigms were assumed to create spherical electric fields in a homogeneous environment and implemented to affect pyramidal neurons within the LIF model based on their distance from the electrode as discussed in the Methods. P1 neurons were uniformly distributed around the electrode from 10*μ*m-2mm away based on staining work by Levitt et al.^31^, and PS pulse parameters by design matched those of Hanks et al.^32^: 10*μ*A, 300*μ*s/phase, 200pulse/s.

In Figure 2, the behavioral effects of PS, CGS, and AGS on decision making were assayed by the percentage of trials in which the stimulated population (P1) won and the decision time. In control trials without electrical stimulation, the decision-making network produced a characteristic psychometric curve with no significant bias (*p* = 0.98 by 1-sample non-parametric bootstrap N=10,000). The percentage of trials in which P1 wins depended on the coherence of the task input, with −100% favoring P2 and +100% favoring P1 (Figure 2A, black).

**Figure 2.**
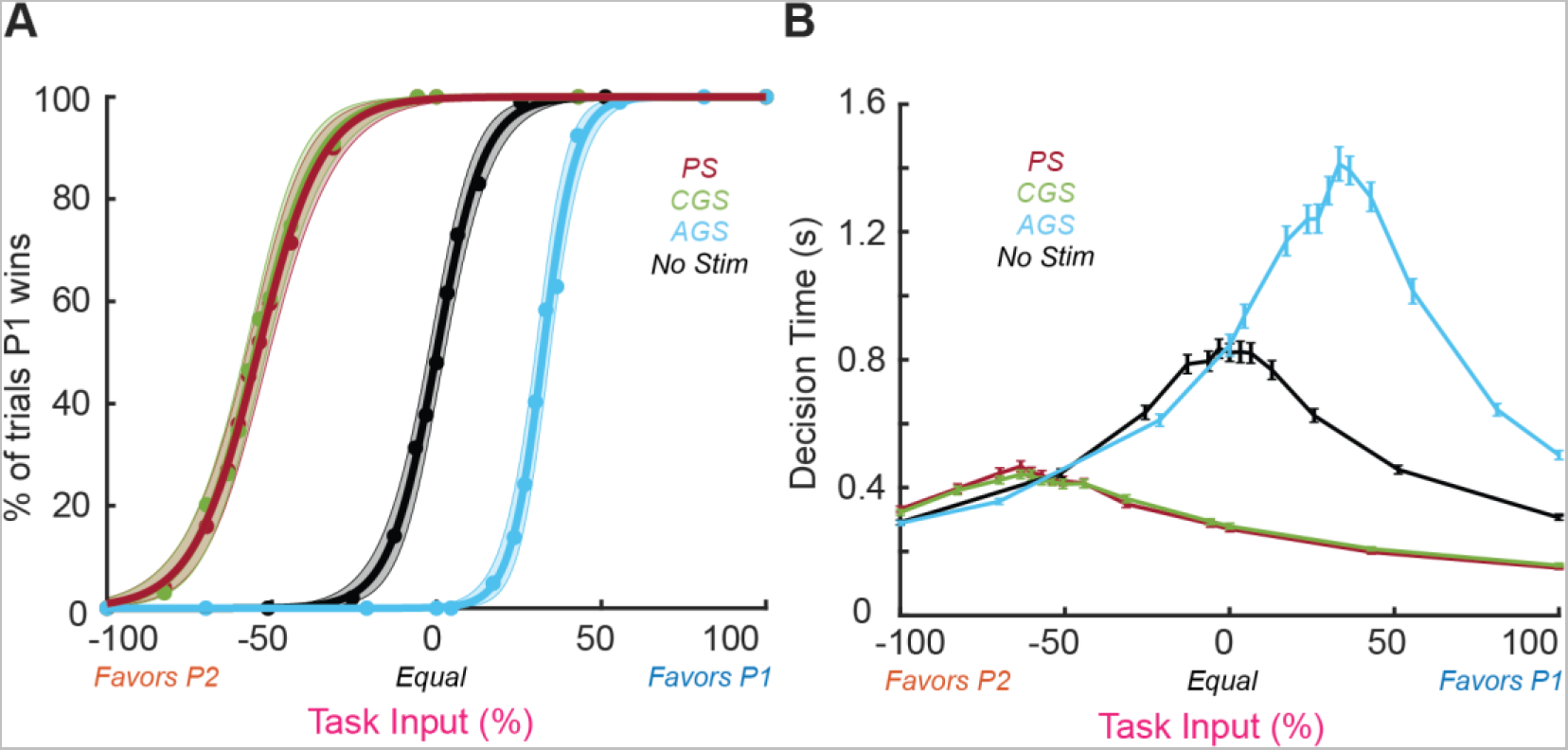
Effects of electrical stimulation on decision making and decision time (N=100 trials). The decision metrics are shown for pulsatile (red) and cathodic galvanic (green), anodic galvanic (blue), and no stimulation (black). (A) The percentage of trials in which the stimulated population (P1) wins the decision-making process are displayed for various levels of visual coherence. Psychometric curves are shown in bold with shaded regions indicating 95% bootstrapped confidence intervals (N=10,000 bootstraps). Red and green curves were calibrated to coincide by design to allow us to compare the subsequent effects of PS and CGS on the neural populations. (B) The time it takes for the winning population to clear the decision threshold (defined to be 15 spk/s) is shown. Error bars depict trial mean and standard error at each coherence level.

To compare the relative effects of excitatory stimulation from PS to those induced by CGS, we calibrated the input from CGS such that the bias of the two stimuli were statistically equivalent. We did this by adjusting the current amplitude of CGS, until its psychometric curve matched that of PS (green and red respectively are equal by design in Figure 2A), resulting in CGS= −1.4µA. We then adopted AGS=−CGS=+1.4µA so that we could make a direct comparison to network behavior based on these stimulation paradigms.

Excitatory stimulation from PS and CGS caused the network to favor P1, shifting the psychometric curve by −54.3%±1.8% and −55.0%±1.7% respectively (Figure 2A; PS in red, CGS in green). Electrical inhibition from AGS caused the network to favor P2, shifting the psychometric curve by +32.3%±1.1% (Figure 2A, blue). Although CGS and AGS used identical amplitude current (1.4*μ*A), AGS shifted the psychometric curve significantly less (*p* < 10^−4^ by 2-sample non-parametric bootstrap N=10,000). CGS and PS induced statistically equivalent shift (*p* = 0.76 by 2-sample non-parametric bootstrap). PS and CGS also broadened the psychometric curve (although only PS was statistically significant), decreasing the slope by 0.94 ±0.24 and 0.83 ± 0.28 respectively relative to no-stimulation (*p* = 0.022, *p* = 0.059 by 2-sample non-parametric bootstrap N=10,000). AGS steepened the psychometric curve, increasing the slope by 1.56 ±0.53 (*p* = 0.009 by 2-sample non-parametric bootstrap N=10,000).

All three stimulation paradigms also affected the decision times (Figure 2B). Excitatory stimulation from PS and CGS decreased peak decision time by 0.53 ±0.02s and 0.58 ±0.02s respectively and shifted the peak decision time by −57.5 ±1.7% and −57.0 ±1.8% (Figure 2B; PS in red, CGS in green). Inhibitory influence from AGS increased peak decision time by 0.69 ±0.04s and shifted the coherence by +31.3 ±1.3% (Figure 2B blue). As with the psychometric curve, AGS shifted the coherence significantly less than CGS (*p* = 3.1 × 10^−23^ by unpaired t-test). However, AGS induced statistically equivalent change in peak decision time as CGS (*p* = 0.34 by unpaired t-test). As with the psychometric curves, CGS and PS induced statistically equivalent coherence shift of the peak decision time (*p* = 0.83 by unpaired t-test).

These changes in decision making elicited by PS in our model are consistent with those observed in behavioral studies^32^, albeit with larger magnitude. Psychometric curves are shifted such that stronger task-related input is required to make decisions against the stimulated population (P1). Decision times are decreased when task-related input and stimulation both favor P1 (e.g. at +25% coherence) but increased when task-related input and stimulation battle over control of the network (e.g. at −60% coherence; Figure 2B, red). CGS showed similar interactions (Figure 2B, green), while the effects of AGS were smaller in magnitude and opposite in polarity (Figure 2B, blue). The decreased effectiveness of AGS relative to CGS was likely due to a floor effect, since neurons cannot decrease their firing rates below 0spk/s from a baseline of only 0-4spk/s. Importantly, both excitatory stimulation paradigms (PS and CGS) pushed P1 firing rates toward the decision threshold and thereby decreased overall decision times; whereas electrical inhibition (AGS) pulled P1 firing rates away from the decision threshold and thereby increased overall decision times. Importantly, AGS inhibited P1 decision times so much that a substantial fraction of trials did not reach the decision threshold before the task period ended at t=3s (Supplementary Figure 3). This effect is consistent with experimental studies using tDCS to bias decision making^33^. Similarly, both excitatory stimulation paradigms (PS and CGS) reduced the slope of the psychometric curves, whereas electrical inhibition (AGS) steepened the slope. These results agree with findings by Salzman et al.^34^ that pulsatile stimulation of area MT significantly flattened psychometric curves. This line of evidence suggests that CGS can effectively mimic PS in the context of perceptual decision making, at a relatively low amplitude (−1.4*μ*A).

### PS and GS Induce Different Profiles of Neuronal Activation/Deactivation

PS and GS affect single neurons differently. PS has a strong depolarizing effect on supra-threshold neurons, primarily causing affected neurons to evoke action potentials in response to each pulse. GS on the other hand, smoothly modulates extracellular potentials, making affected neurons more likely or less likely to fire action potentials in response to EPSPs. We hypothesized that the effects of PS on neurons will be less affected by other neural connections due to its strong ability to evoke spikes.

#### Responses in a Completely Disconnected Network

To investigate the mechanism of how PS and GS interact with the neural network, we first ascertained the effects of PS and GS in disconnected neurons, leveraging the benefits of the experiment being conducted entirely *in-silico*. Disconnecting the neurons in P1 allowed us to measure firing rates over the entire task period (t=1-3s) without the confounds induced by network activity (Figure 3A).

**Figure 3.**
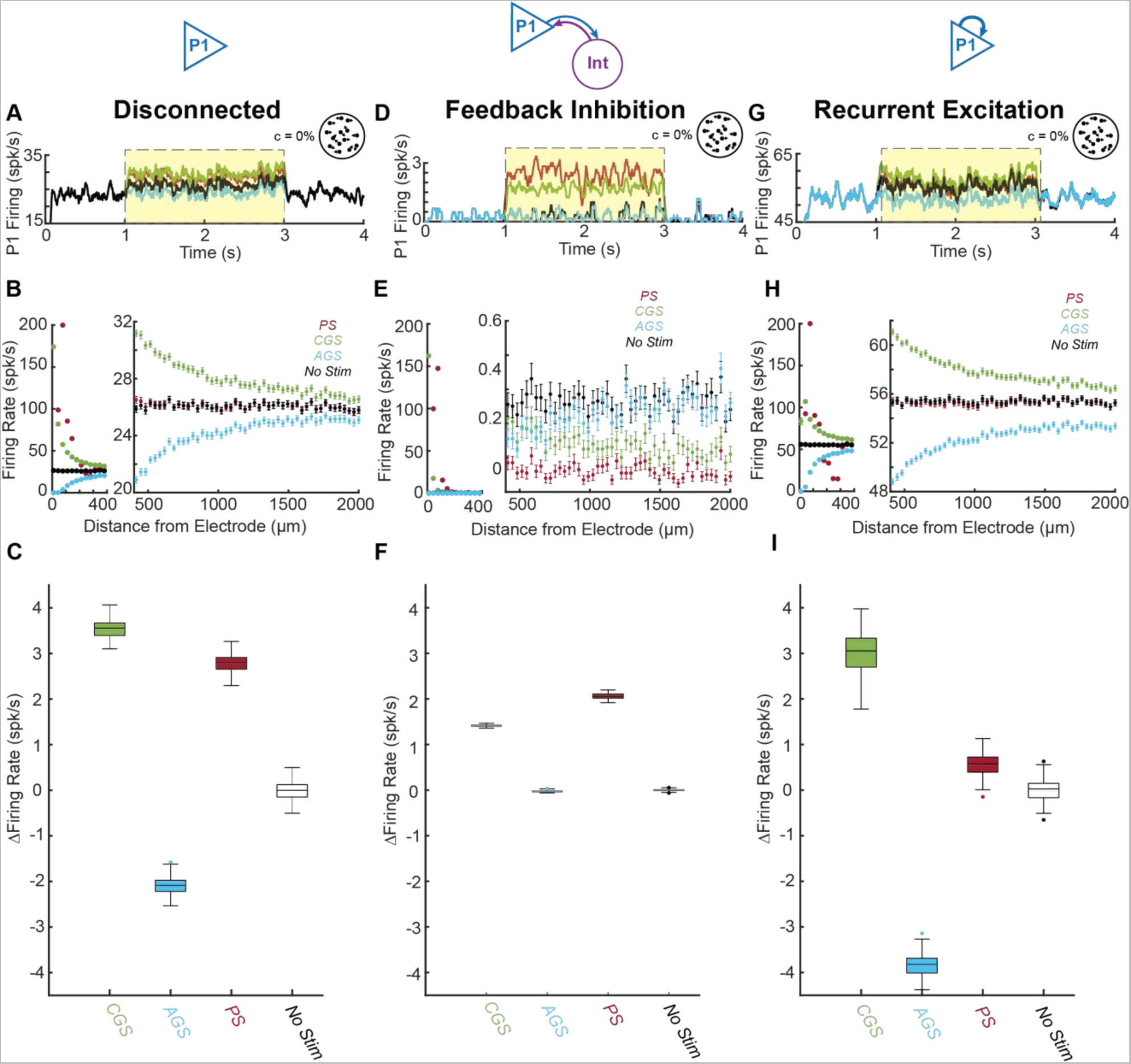
Effects of pulsatile (red), cathodic galvanic (green), anodic galvanic (blue), and control (black) stimulation on individual P1 neural firing rates (N=100 trials). Networks with fully disconnected neurons (A-C), feedback inhibition only (D-F), and recurrent excitation only (G-I) are investigated. Average P1 Firing for representative trials are shown in A, D and G. Each neuron’s mean task firing rate ±sem (t=1-3s) is shown as a function of its distance to the stimulation electrode during stimulation (yellow zone) (B, E, H; left: neurons <400*μ*m from electrode, right; neurons > 400*μ*m from electrode). Box plots (C, F, I) depict each trial’s population-averaged change in firing rate relative to no stimulation. For all trials, task-related input was equal for P1 and P2 (coherence = 0%).

As expected from single neuron studies, during stimulation (Figure 3A, yellow zone) the distributions of firing rates induced by PS and CGS were significantly different from each other and matched the expectations of single neural responses to stimulation (*p* = 1.1 × 10^−18^ by Kolmogorov-Smirnov test; Figure 3B).

For PS, due to the refractory effects of high-amplitude pulses, the neurons closest to the pulsatile stimulation electrode (<40*μ*m) were blocked. Most of the neurons farther away (44-314*μ*m) were excited with decreasing level of excitation as a function of distance. Excitation was limited to the neurons close to the pulsatile stimulation electrode up to 347*μ*m from the stimulation site (*p* > 0.05 by unpaired 1-tailed t-test with Bonferroni correction; Figure 3B red, right panel).

In contrast, for CGS, the neurons closest to the stimulation electrode (<400*μ*m) were strongly excited with firing rates up to 173spk/s (Figure 3B green, left panel). In addition, weak excitation spread far from the galvanic stimulation electrode with small but significant increases in neural firing rates up to 1798*μ*m away (*p* < 0.05 by unpaired 1-tailed t-test with Bonferroni correction; Figure 3B green, right).

Compared to CGS, AGS had equal and opposite effects on neural firing rates for neurons >246*μm* away (*p* > 0.05 by unpaired 2-tailed t-test with Bonferroni correction; Figure 3B, blue). However, it completely blocked the activity of neurons <50*μm* away and thereby induced a smaller change in firing rate compared to CGS (Figure 3B, blue). As a result, AGS induced a smaller average change in firing rate (median: 2.09spk/s IQR: 1.97-2.22spk/s) compared to CGS (median: 3.56spk/s IQR: 3.39-3.67spk/s) in the P1 population (*p* = 2.6 × 10^−34^ by Kruskal-Wallis test; Figure 3C).

PS induced a smaller change in firing rate (median: 2.80spk/s IQR: 2.65-2.91spk/s) than CGS when averaged over the entire neural population of P1, likely due to its smaller spread of activation (*p* = 6.7 × 10^−9^ by Kruskal-Wallis test; Figure 3C).

From these results in disconnected neurons, one would expect that CGS would induce a greater change in decision-making than PS since it affects more neurons and induces a greater overall change in firing rate. However, by design they induce identical effects in a fully connected network (Figure 2A), suggesting that PS and GS interact with the interconnected network in more complicated ways.

#### Network with Only Inhibitory Feedback

We next probed the effects PS and GS in the context of feedback inhibition. We added 30 inhibitory interneurons connected to the P1 population to create the 80/20 E/I ratio, which yielded a stable P1 spiking activity during PS and CGS (Figure 3D, yellow zone).

Feedback inhibition dramatically reduced spontaneous firing rates from ~25spk/s to <1spk/s. Nevertheless, some neurons closest to the CGS and PS electrodes (<100*μ*m) were still strongly activated achieving firing rates up to 162spk/s from CGS and 147spk/s from PS (Figure 3E, left panel). However, the far-reaching weak excitation induced by CGS seen in the disconnected network (Figure 3B, right panel) was severely attenuated by compensatory feedback inhibition (Figure 3E, right panel). CGS only induced increases in neural firing rates up to 178*μm* away from the electrode (*p* < 0.05 by unpaired 1-tailed t-test with Bonferroni correction; Figure 3E green, right panel).

P1 neurons stimulated by PS also experienced compensatory feedback inhibition, causing 51 neurons 500-2000*μ*m away from the electrode to have lower firing rates than non-stimulated control (*p* < 0.05 by unpaired 1-tailed t-test with Bonferroni correction; Figure 3E red, right panel).

As a result, both CGS and PS were significantly less effective at increasing neural firing rates under feedback inhibition than with disconnected neurons (*p* = 2.6 × 10^−34^ and *p* = 2.6 × 10^−34^ respectively by Wilcoxon rank sum test; Figure 3C&F). Importantly, however, CGS induced a smaller average increase in firing rate (median: 1.41spk/s IQR: 1.40-1.43) than PS (median: 2.06spk/s IQR: 2.02-2.11spk/s) under feedback inhibition (*p* = 5.6 × 10^−9^ by Kruskal-Wallis test), indicating that the effects of PS are more resistant to network-based suppression.

The effects of AGS were almost completely indistinguishable from the non-stimulated control under strong feedback inhibition (Figure 3E&F), providing further support for the hypothesis that the limited effectiveness of AGS is driven by a floor effect.

From these results in neurons exposed to feedback inhibition only, one would expect that PS would induce a greater change in decision making than CGS since it induces a greater increase in average firing rate. On the contrary, we observed an equivalent effect, indicating that another network effect may be influencing decisions.

#### Network with Only Excitatory Feedback

Finally, we investigated the effects of GS and PS with recurrently connected P1 neurons only. To avoid runaway recurrent excitation, we added *weak* recurrent excitatory connections (*w_rec_*=0.05) among the neurons in P1 and still maintained a stationary process (Figure 3G). Recurrent excitation dramatically increased spontaneous firing rates from ~25spk/s to ~52spk/s.

Despite the increased baseline firing rates, the neurons exposed to PS experienced a similar activation profile as in the disconnected case with a few neurons firing ~90spk/s, one neuron fully activated at 200spk/s, and neurons >145*μ*m away not significantly increased relative to control (*p* < 0.05 by unpaired 1-tailed t-test with Bonferroni correction; Figure 3H, red).

In contrast, CGS activation scaled synergistically with the recurrent excitation, inducing higher firing rates than in the disconnected case for all but the closest neuron (Figure 3H, green; compared to Figure 3B, green). Due to the high excitatory current from both CGS and recurrent inputs, the closest neuron experienced intermittent depolarizing block, which caused its average firing rate to decrease to 82±7spk/s, which was highly variable among trials. Importantly, CGS induced a significantly greater increase in firing rate than PS under recurrent excitation (*p* = 7.6 × 10^−10^ by Kruskal-Wallis test), indicating that the effects of CGS are more synergistic with network-based excitation.

Under recurrent excitation, AGS induced a larger change in firing rates (median: 3.82spk/s IQR: 3.69-4.01spk/s) than in the disconnected case (median: 2.09spk/s IQR: 1.97-2.22spk/s) due to the elevated spontaneous firing rates and removed floor effect (*p* = 2.6 × 10^−34^ by Kruskal-Wallis test; Figure 3C&I, blue).

Since membrane voltage changes in response to suprathreshold pulses are not affected by small fluctuations in membrane potential, neurons that are directly affected by PS are not very sensitive to either feedback inhibition or recurrent excitation. In contrast, GS induces small changes in membrane potential that increase (for CGS) or decrease (for AGS) the probability of firing action potentials in response to naturally occurring EPSCs and IPSCs^1^^9^. For this reason, CGS and AGS are more sensitive to the ongoing network activity, and significantly alter average firing rates farther from the site of the electrode than PS. As a result of these phenomena, increasing the strength of feedback inhibition in the network gives PS more influence on the network activity relative to GS; whereas increasing the strength of recurrent excitation in the network gives GS more influence over network activity relative to PS.

### PS vs. GS Effectiveness Depends on Dynamic E/I Balance Changes

Based on the results obtained with modified excitation/inhibition in the previous section, we hypothesized that the ability of GS to modulate the fully connected network activity would be highly dependent on the excitatory/inhibitory balance. In contrast, the ability of PS to evoke spikes should largely remain unchanged independent of this network behavior.

Because of the recurrent nature of the network, the relative strengths of feedback inhibition and recurrent excitation change during the time course of each trial, with or without electrical stimulation. We measured the relative impact of these two network motifs by recording the recurrent AMPA, NMDA, and GABA currents experienced by each neuron in P1. We indexed the overall *network current* by adding the recurrent excitatory AMPA and NMDA currents and subtracting the inhibitory GABA current. Thus, when the *network current* is negative, the network favors feedback inhibition, but when it becomes positive it favors recurrent excitation (Figure 4A&B).

**Figure 4.**
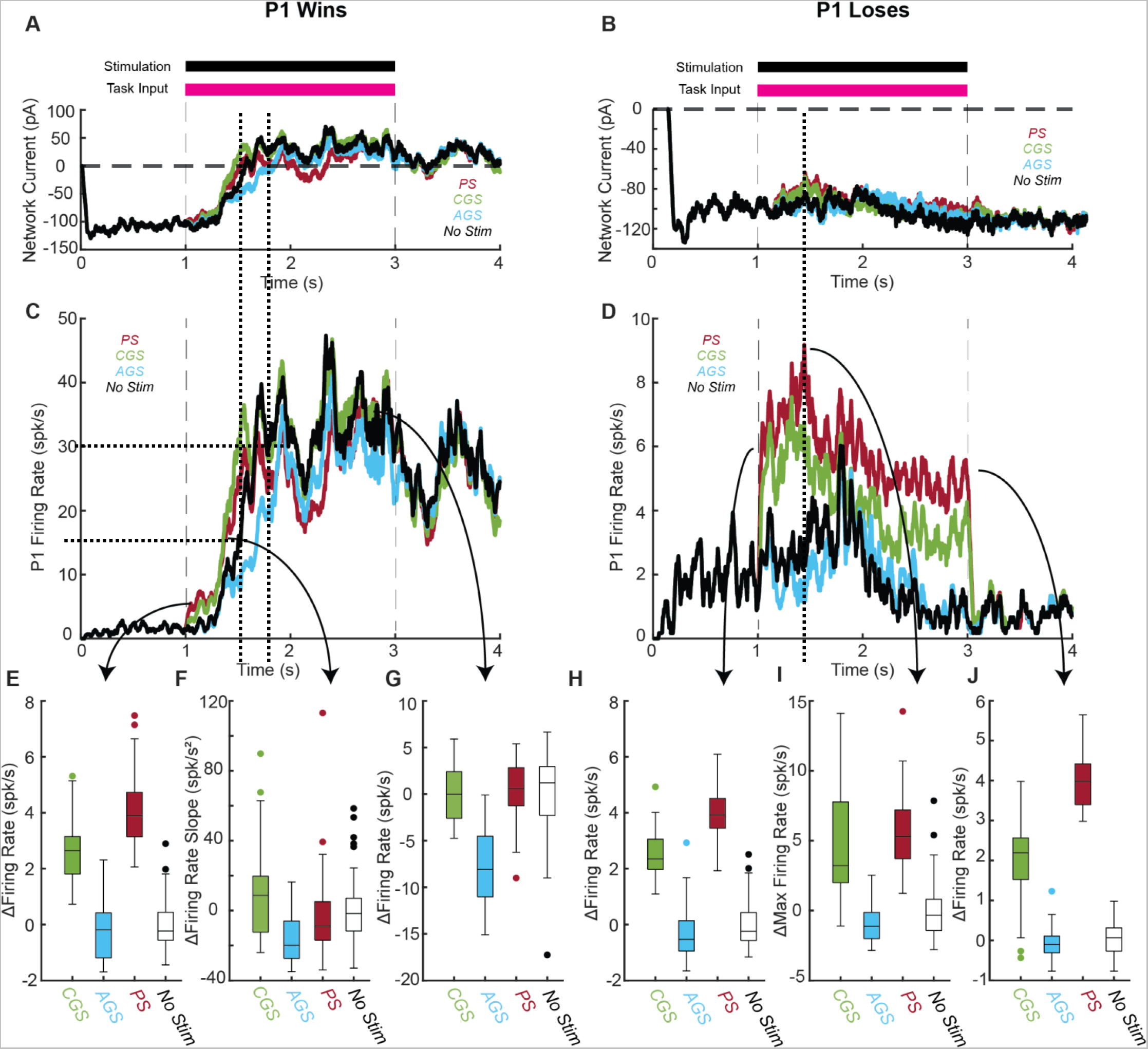
Effects of pulsatile (red), cathodic galvanic (green), anodic galvanic (blue), and control (black) stimulation on P1 population firing rates over time (N=100 trials). Trials in which P1 won (left, A, C, E-G) and P1 lost (right, B, D, H-J) are investigated separately. (A, B) Network currents (*I_AMPArec_* + *I_NMDA_* − *I_GABA_*) received by P1 neurons are shown in two representative trials. (C, D) P1 firing rate trajectories are shown over time in two representative trials. (E, H) Box plots depict the population-averaged change in firing rate relative to no stimulation for the first 100ms of the trial (t=1-1.1s). (F) Box plots depict the population-averaged change in firing rate slope around the decision threshold of 15spk/s relative to No Stim. (I) Box plots depict the population-averaged maximum firing rate relative to No Stim. (G, J) Box plots depict the population-averaged change in firing rate relative to No Stim for the last 100ms of the trial (t=2.9-3s). For all trials, task-related input was set such that P1 and P2 each won 50% of the trials (coherence = 0% for No Stim, −57% for Pulsatile and Cathodic Galvanic, and +30% for Anodic Galvanic). Only trials in which P1 won before t=2.5s were included in the analysis.

At baseline, feedback inhibition is substantially stronger than recurrent excitation (≈ −100pA), which holds the pyramidal neuron firing rates at 0-4spk/s (Figure 4A-D). Once the task-related input turns on at t=1s, input excitation and recurrent excitation together overpower feedback inhibition and pyramidal firing rates begin to rise. As P1 firing rates rise, recurrent excitation grows, creating a positive feedback loop. If P1 wins, when average P1 firing rates clear the critical decision threshold of 15spk/s, this positive feedback loop overwhelms feedback inhibition, and P1 firing rates surge rapidly up to ~30spk/s. In this elevated firing rate state, recurrent excitation is dominant (+25-50pA) and sustains high firing rates (30-40spk/s) even after the task-related input turns off at t=3s (Figure 4A&C dotted lines identify time indices for 15 and 30 spk/s). If P1 loses, it is instead strongly suppressed by feedforward inhibition after P2 clears the decision-making threshold. In this case, feedback inhibition is dominant throughout the trial, but recurrent excitation is most competitive (≈ −70*pA*) just before P2 clears the decision-making threshold, when both populations’ firing rates are elevated (~10spk/s) (Figure 4B&D, dotted line). Therefore, the relative strengths of feedback inhibition and recurrent excitation depend on both the outcome (P1 winning or losing) and time course of each trial.

Based on the results in the simplified networks, we hypothesized that PS would become more effective as feedback inhibition became more dominant, and CGS would become more effective as recurrent excitation became more dominant in the network. These differences should be apparent in the firing rate trajectories of winning and losing trials. Indeed, we observed distinct differences in the average P1 firing rate trajectories between PS, CGS, and AGS (Figure 4C-J). These differences occurred even though PS and CGS induced equivalent changes in decision making.

We first assessed the effectiveness of PS and GS at the beginning of each trial to understand their immediate effects on the decision-making network (Figure 4E, H). In the first 100ms of the task period (t=1-1.1s) PS caused a larger increase in P1 firing rates (median: 3.89spk/s IQR: 3.14-4.73spk/s) compared to CGS (median: 2.64spk/s IQR: 1.81-3.14spk/s) in trials in which P1 won (Figure 4E; *p* = 0.078 by Kruskal-Wallis test). PS also caused a significantly larger increase in P1 firing rates (median: 3.93spk/s IQR: 3.45-4.51spk/s) than CGS (median: 2.34spk/s IQR: 1.97-3.05spk/s) in the first 100ms (t=1-1.1s) of trials in which P1 lost (Figure 4H; *p* = 6.9 × 10^−3^ by Kruskal-Wallis test). This result is consistent with the hypothesis that PS is more effective than CGS when feedback inhibition is strongly dominant in the network (≈ −100pA at 1.1s in Figure 4A,B). This finding is independent of the outcome of the trial because feedback inhibition is dominant at the beginning of both winning and losing trials.

Next, we assessed the effectiveness of PS and GS around the decision-making threshold (Figure 4F, I) to understand how their effects integrate with dynamic network activity.

In winning trials, we observed that the slope of the P1 firing rate curve around the decision threshold (15spk/s) was significantly affected by both PS and CGS (Figure 4F; *p* = 4.2 × 10^−5^ by Kruskal-Wallis test). The slope of the firing rate was increased by CGS (median: 8.72spk/s^2 IQR: −12.38-19.63spk/s^2), but decreased by PS (median: −8.89spk/s^2 IQR: −17.01-5.08spk/s^2) and AGS (median: −19.82spk/s^2 IQR: −27.31-(−6.00)spk/s^2) compared to No Stim (Figure 4F). As a result, CGS had a significantly higher median slope than PS (*p* = 0.03 by Kruskal-Wallis test) and AGS (*p* = 0.005 by Kruskal-Wallis test). This finding is consistent with the hypothesis that CGS is synergistic with recurrent excitation, since recurrent excitation drives the rapid increase in P1 firing rates around the decision threshold of 15spk/s. On the other hand, PS-induced activation does not synergize well with recurrent excitation, limiting its effectiveness to rapidly increase firing rates around the decision threshold. The inhibitory effectiveness of AGS is increased as P1 firing rates increase due to the relaxing of the floor effect, which also results in a shallower slope.

In losing trials, we observed that the maximum P1 firing rate was increased by PS (median: 5.30spk/s, IQR: 3.72-7.20spk/s) and CGS (median: 3.21, IQR: 1.99-7.78spk/s), but decreased by AGS (median: −1.13spk/s, IQR: −2.01-(−0.14)spk/s) relative to No Stim (Figure 4D, I). PS and CGS induced statistically equivalent increases in maximum P1 firing rate (*p* = 0.44 by Kruskal-Wallis test). Our explanation for these equivalent increases in firing rate is that CGS becomes more effective relative to PS with even a slight increase in recurrent excitation (Figure 4D, dotted line).

Finally, we assessed the effectiveness of PS and GS at the end of each trial to understand their long-term effects on the network in steady state (Figure 4G, J). In the last 100ms of the task period (t=2.9-3s), average P1 firing rates were increased by PS (median: 0.58spk/s IQR: −1.25-+2.83spk/s), not affected by CGS (median: 0.00spk/s IQR: −2.59-+2.41spk/s), and decreased by AGS (median: −8.09spk/s IQR: (−11.02)-(−4.52)) relative to No Stim in trials in which P1 won (Figure 4G). Crucially, PS and CGS induced statistically equivalent changes in P1 firing rates relative to No Stim (*p* = 0.82 by Kruskal-Wallis test). This finding was somewhat surprising, and not immediately consistent with our hypothesis. Recurrent excitation is strongly dominant at the ends of trials in which P1 wins (+25-50pA at 3s; Figure 4A), so we expected CGS to induce a greater increase in P1 firing rates than PS. Instead, we observed that CGS was equivalent to and even slightly less effective than PS. Upon further investigation, we found that shortly after P1 wins, the dramatic increase in recurrent excitation caused one of the P1 neurons to experience depolarizing block from CGS. This outlier drastically reduced average P1 firing rates, while not affecting the decision-making outcome of the trial, since the decision threshold had already been cleared.

In the last 100ms of losing trials (t=2.9-3s), P1 firing rates were increased by PS (median: 3.98spk/s IQR: 3.40-4.42spk/s) and CGS (median: 2.19spk/s IQR: 1.52-2.56spk/s) but decreased by AGS (median: −0.10spk/s IQR: −0.31-(+0.11)spk/s) relative to no stimulation (Figure 4J). As expected, PS caused a significantly larger increase in P1 firing rates than CGS (*p* = 3.8 × 10^−4^ by Kruskal-Wallis test). These findings are consistent with the hypothesis that increased feedback inhibition favors PS relative to CGS. At the ends of trials in which P1 loses, feedback inhibition is as dominant as it gets (−100-(−120)pA), and the difference between PS and CGS is also at its maximum (~2spk/s). Additionally, AGS is much more effective in winning trials than losing trials due to the relief of the floor effect. These differences in P1 firing rate trajectories over the time course of each trial represent measurable predictions about the different effects of PS and GS on functional networks of neurons that arise directly from a mechanistic understanding of their differing responses to network motifs of excitation and inhibition.

### PS and GS Induce Different Spatiotemporal Distributions of Activation

Because PS appears to be generally less susceptible to network effects, we expect to find high variability in neural responses as we examine the PS effect on neurons at different distances from the electrode. PS should affect neurons nearby more than neurons farther away. In contrast, we would expect the effect of GS to be more uniform and less dependent on distance. To examine these hypotheses, we investigated the distribution of neural firing rates in the fully connected network. We visualized the neural activity as it evolved during the time course of each trial and as a function of distance from the electrode. To quantify the differences in variability of neural firing rates across the population of neurons, we used the fourth central moment: kurtosis as a measure of variability of firing rates across the neural population (Figure 5B). High kurtosis values indicate high variability and low kurtosis values indicate uniformity of firing rates across population. We selected this metric due to its ability to highlight the most dramatic changes in neural firing rates such as blocking (<1spk/s) and hyper-excitation (>100spk/s).

**Figure 5.**
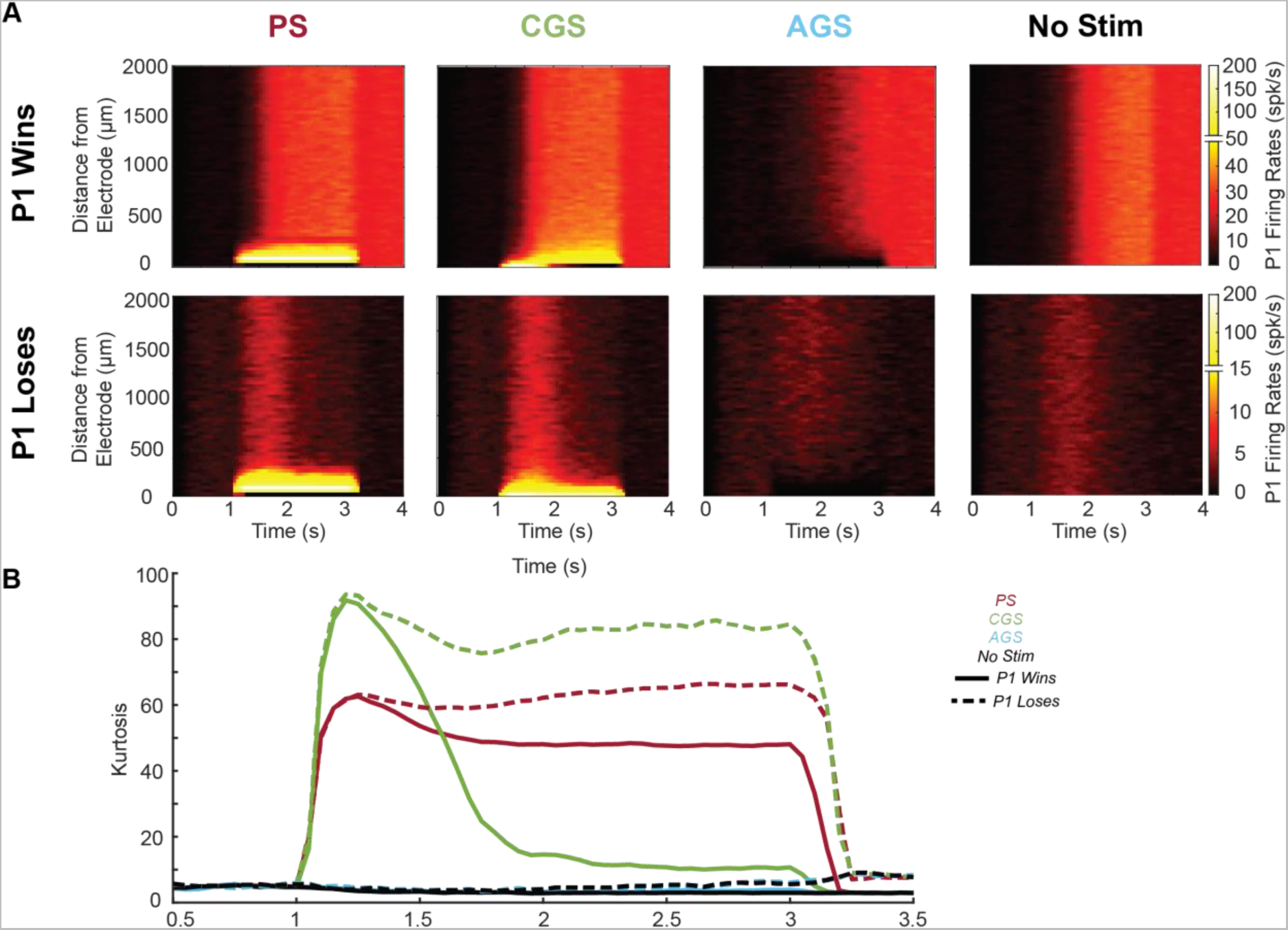
Effects of pulsatile (red), cathodic galvanic (green), anodic galvanic (blue), and control (black) stimulation on distributions of P1 firing rates over time (N=100 trials). (A) Heatmaps show individual neural firing rates for trials in which P1won (top) trials in which P1 lost (bottom). (B) Kurtosis of the distribution of P1 firing rates is shown over time. Trials in which P1 won are represented by solid lines, and trials in which P1 lost are shown by dashed lines. Beginning (0-0.5s) and end (3.5-4s) of trials are excluded due to artifactual effects when population firing rates are too low (<1spk/s). For all trials, task-related input was set such that P1 and P2 each won 50% of the trials (coherence = 0% for No Stim, −57% for Pulsatile and Cathodic Galvanic, and +30% for Anodic Galvanic). Only trials in which P1 won or lost before t=3s were included.

As expected from the disconnected trials, CGS and PS strongly excited a small subset of neurons close to the electrode (<500*μ*m) up to 200spk/s (Figure 5A). PS also induced full pulse-pulse blocking in the closest neuron. These strong local effects are seen as increases in kurtosis at the beginning of the task period (t=1.1-1.2s) of both winning and losing trials (Figure 5B). During this onset, CGS strongly activated only the closest neuron, whereas PS activated a larger block of close by neurons. As a result, CGS induced significantly larger kurtosis transients (median: 82.87 IQR: 76.63-87.83) compared to PS (median: 57.31 IQR: 53.51-60.48 *p* = 2.6 × 10^−5^ by Kruskal-Wallis test).

In winning trials, PS caused relatively static excitation (Figure 5B, solid red), but CGS excitation changed with time (Figure 5B, solid green). As P1 firing rates increased, CGS activation spread due to the increased recurrent excitation, captured by the falling kurtosis levels. PS activation also spread, but much less than CGS. As a result, by the end of the task period (t=2.9-3s) of winning trials, PS maintained the large nonuniformity in firing rates, shown by the significantly higher kurtosis (median: 47.88 IQR: 46.56-49.53) than CGS (median: 10.48 IQR: 9.19-11.51 *p* = 2.4 × 10^−5^).

In losing trials, P1 firing rates achieved a maximum around 1.5s before P2 cleared the decision-making threshold, after which P1 was suppressed. Accordingly, CGS achieved its maximal spread throughout the network during this time, and then reverted to strongly activating only a few neurons (Figure 5A, bottom). As a result, similar to the initial transients, CGS maintained higher (but not significantly higher) kurtosis (median: 101.61 IQR: 48.96-106.15; Figure 5B, dashed green) than PS (median: 66.43 IQR: 64.70-68.71; Figure 5B, dashed red) at the end of the task period (t=2.9-3s) of trials in which P1 lost (*p* = 0.22 by Kruskal-Wallis test).

AGS induced a mild deactivation, especially prominent for the closest neurons (<500*μ*m) in winning trials. AGS did not affect firing rates dramatically in losing trials due to the floor effect. As a result, AGS did not induce significant changes in firing rate distributions compared to No Stim (*p* > 0.05 by Kruskal-Wallis test for all comparisons).

These findings support the hypothesis that CGS induces a more uniform spread of activation that is highly network-dependent, whereas PS activates a single block of neurons relatively statically throughout the task period. Interestingly, at the beginning of the task period, when P1 firing rates are low, CGS induced higher kurtosis than PS (Figure 5B). This indicates that CGS takes some time for its effects to spread throughout the network, and initially strongly activates fewer neurons than PS. Similarly, when P1 lost and firing rates remained low, CGS induced higher kurtosis than PS, suggesting that high levels of recurrent excitation relative to feedback inhibition are necessary to propagate the effects of CGS throughout the network. This finding is consistent with the results from the simplified networks, in which feedback inhibition attenuated the spread of activation induced by CGS.

### PS but not GS Induces Synchronous Firing in the Closest Neurons

The decision-making model employed here assumes that spatiotemporal integration of neural firing exclusively determines the perceptual decision-making process. However, recent evidence suggests that precise spike timing may play a complementary role in determining when to prioritize certain streams of information over others^35,36^. Given that PS is more likely to influence neurons independent of network activity, we expect that the effect of PS on neural firing would be stronger and more phase-locked at the start of the stimulation, while the effect of GS on the population would be more distributed and less correlated.

To understand the steady-state effects of each stimulation modality, we measured spike timing in the last 0.5s of the task period (t=2.5-3s; Figure 6A, yellow). We found that PS induced fully phase-locked activity in the responding neurons closest to the electrode (<300*μ*m), and partially phase-locked activity (20-60% phase-locked APs) in neurons a moderate distance away (300-400*μ*m) in P1 (Figure 6B, red). For each neuron in P1, we computed the coefficient of variation (CV) to measure regularity of firing. PS induced highly regular firing (0-0.4) in a small number of neurons closest to the electrode (<150*μ*m); whereas CGS and AGS caused a more modest increase in regularity (Figure 6C). Last, for each neuron pair in P1, we calculated the percentage of synchronized APs (<300*μ*s apart) in Figure 6D. We observed a large increase in synchrony among the neurons phase-locked to pulses (up to 100% synchronized <300*μ*m from the electrode; Figure 6D, left). These neurons were the primary drivers of an overall increase in synchrony from PS (*p* < 10^−15^ by 1-way ANOVA). In contrast, CGS did not induce significant synchrony compared to control (Figure 6D, middle-left; *p* =0.93 by 1-way ANOVA). AGS caused a mild desynchronizing effect (Figure 6D, middle-right; *p* < 10^−15^ by 1-way ANOVA).

**Figure 6.**
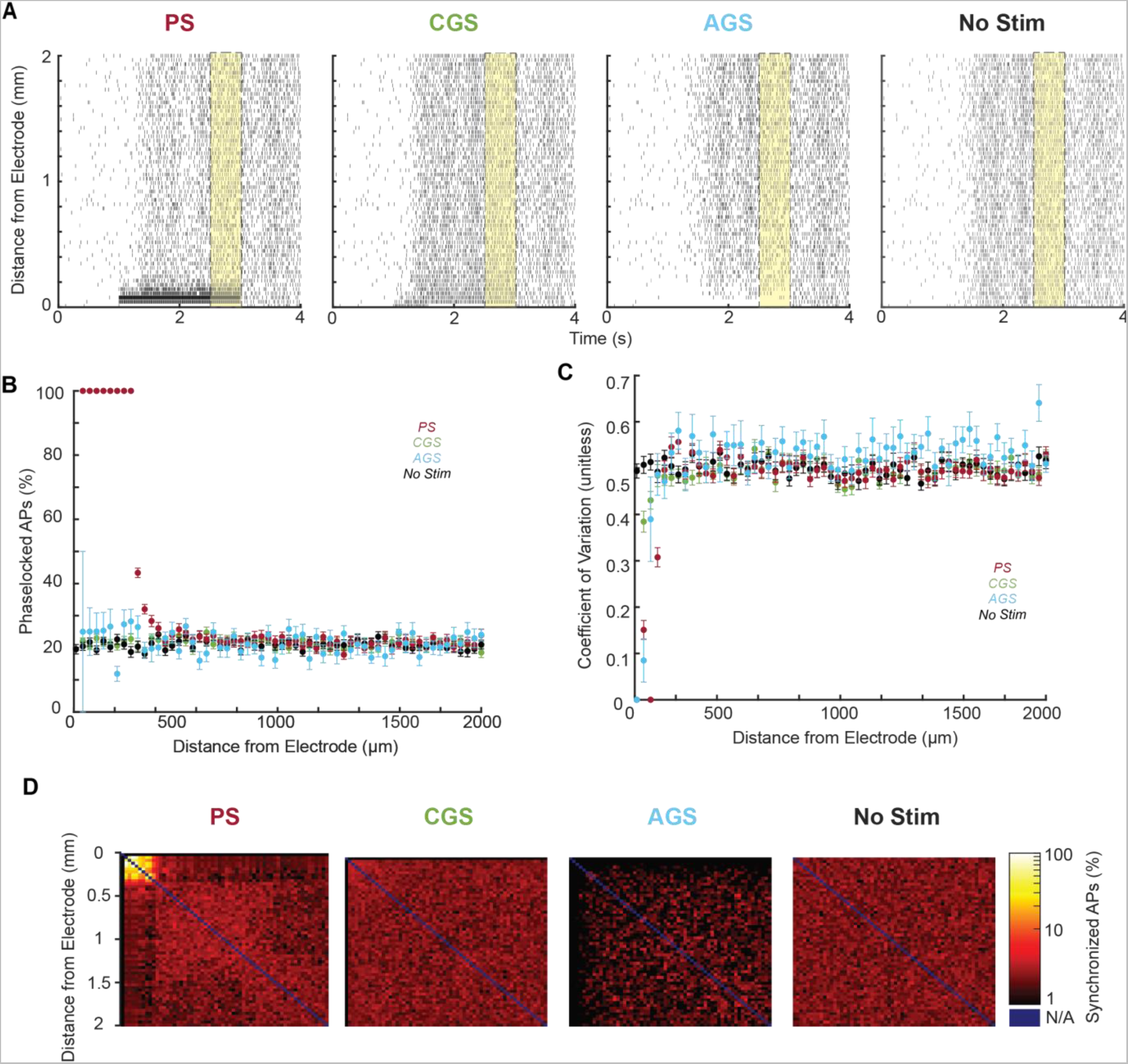
Spike timing differences in P1 neurons among stimulation conditions (Pulsatile, red; Cathodic Galvanic, green; Anodic Galvanic, blue; and No Stimulation, black) for N=100 trials. (A) Representative trial rasters are plotted for each stimulation condition, with end-of-task period (t=2.5-3s) highlighted in yellow. (B) The percent of each neuron’s action potentials that occur during a pulse presentation for the end-of-task period is shown as a function of its distance to the stimulation electrode. (C) Each neuron’s end-of-task coefficient of variation (CV) is shown as a function of its distance to the stimulation electrode. (D) Heat map of the percent of action potentials from a neuron that are synchronized to another neuron as a function of distance from the stimulation electrode. Self-synchrony was undefined (N/A, blue). For all trials, task-related input was set such that P1 and P2 each won 50% of the trials (coherence = 0% for No Stim, −57% for Pulsatile and Cathodic Galvanic, and +30% for Anodic Galvanic). Only trials in which P1 won before t=3s were included.

From these results, we conclude that PS induces phase-locked APs in the directly activated population, which manifest synchronized connections in that subset of neurons. CGS largely preserves spike timing and AGS induces a mild desynchronization. Interestingly, based on the CV statistic, these neurons are not firing as regularly as one might expect. This is likely due to neurons phase-locking to an irregular subset of pulses based on variable relative refractory periods.

## IV. DISCUSSION

By design, both CGS and PS achieved the same strong biasing effect and decreased decision time. Given this equivalence, the two stimulation methodologies exhibited nuanced differences in the ways that they interacted with the decision-making network to achieve this bias. PS elicited responses within the network by strongly and synchronously activating a static block of neurons close to the electrode during both winning and losing trials. In contrast, CGS directly activated a much smaller number of neurons, which yielded a smaller increase in firing rates at the task onset. As the task progressed, the synergy between CGS and recurrent excitation led to faster accumulation of perceptual evidence and a steeper slope in the transition between low P1 firing and high P1 firing. This synergy also contributed to a broader spread of activation in winning trials. When P1 lost, CGS’s sensitivity to feedback inhibition allowed P1 to be appropriately suppressed, but with some unnatural residual activation. Throughout winning trials, CGS did not induce any additional synchrony, generally respecting the spike timing transmitted from the EPSCs/IPSCs analogous to that of the non-stimulated trials.

One limitation of this study is that the effect of PS on decision making in the model is larger than the effect *in vivo* with the same stimulation parameters^32^. Similar studies with lower pulse amplitude (5 *μ*A) were shown to achieve an equivalent effect on decision making in some experiments, also highlighting the apparent inter-subject variability^7,37^. Another limitation is that we only considered electrical activation of excitatory pyramidal neurons in our model, even though inhibitory neurons are co-located in the decision-making circuit. Inhibitory neurons have much smaller axons, so they experience a much weaker effect of electrical stimulation. Also, similar modeling studies suggest that the effects of electrical stimulation on decision-making are primarily driven by its effect on excitatory pyramidal neurons rather than inhibitory interneurons^33^. Nevertheless, there may be some nuanced effect via the interneurons that would be interesting to explore in future work. Finally, we only assessed one pair of PS and GS amplitudes, each of which gives rise to a specific pattern of activation/deactivation depending on distance from the electrode. We chose this approach to enable direct comparison to existing experimental work in the literature^32^, but exploring more equivalent amplitude pairs would help confirm or refute the generalizability our findings.

Based on the modeling experiments in this work, galvanic stimulation appears able to interact with functional neural circuits as effectively as pulsatile stimulation, while minimizing undesired side-effects such as excessive neural synchrony, uneven distributions of neural activation, and insensitivity to ongoing network dynamics. These effects were largely consistent with our hypotheses based on previous modeling work in single neurons but differed in small ways idiosyncratic to the stimulation protocol we used. Specifically, our CGS amplitude was sufficient to directly activate a few neurons closest to the electrode regardless of EPSCs/IPSCs. These neurons fired more regularly (as measured by CV), remained somewhat active under feedback inhibition, and one neuron experienced depolarizing block under sufficient recurrent excitation. This small minority differed from the overall trends of CGS activation and was responsible for unexpected findings such as the decrease in efficacy at the ends of trials in which P1 won. Future work exploring different parameter combinations of PS and CGS will be instrumental in determining whether these depolarizing block effects consistently arise or depend on specific CGS amplitudes.

In addition to excitation, galvanic stimulation can readily support electrical inhibition via AGS. Inhibiting P1 neurons via AGS caused a behavioral bias opposite in polarity, but smaller in magnitude than the bias induced by CGS, due to the floor effect on firing rates. Inhibitory AGS also increased decision times and psychometric sensitivity, whereas excitatory CGS and PS decreased decision times and psychometric sensitivity. These findings, together with related work^3,33,34^, paint a picture in which excitation drives fast, imprecise decisions, and inhibition drives slow, precise decisions. Despite its substantial behavioral bias, AGS did not induce any alterations in firing rate distributions, and it avoids the risks of excessive neural activation such as excitotoxic shock. Therefore, AGS may be an even better candidate for effective neural interfacing than CGS. Feedback inhibition and recurrent excitation are crucial components of the decision-making network, and disruptions to their operation have been shown to impair the decision-making process^3^. Similarly, synchrony effects are important to a variety of cognitive processes^35^ and disease states such as epilepsy^36^. Furthermore, population firing rate distributions follow a characteristic long-tailed structure throughout the brain thought to support sparse neural coding of information^38^. Therefore, although electrical pulses are clearly an effective way to alter decision-making circuits and, more broadly, to interface with the nervous system at large, they may struggle to replicate nuanced interactions that depend on precise spike timing, on-going network activity, and neuron-specific coding. GS generally preserves neural spike-timing, respects on-going network activity, and maintains population firing rate distributions.

The future of effective cortical neuromodulation technology that would replicate normal function may lie in using the combination of the two methods. While GS can bias the network in a more natural way, PS can be used to deliver more tightly localized and precise neural responses.

## VI. STAR METHODS

### Perceptual Decision-making Task

A random dot motion task was simulated at coherence levels from fully leftward (+100%) to fully rightward (−100%) coherence (Figure 1A, circles). 100 trials were executed at each coherence level under four conditions: pulsatile (PS), cathodic galvanic (CGS), anodic galvanic (AGS), and no-stimulation control. Throughout each 4-second trial, all neurons received 2400 Hz background Poisson inputs that triggered AMPA EPSCs. This caused the pyramidal neurons in the network to fire spontaneously at 2-3 spk/s (Figure 1B). At *t* = 1s, neurons in populations P1 (blue) and P2 (red) received task-related input proportional to coherence (*c*):

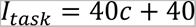

where if motion is in the opposite direction *c* is negative^2^ (Figure 1A-B, magenta). To bias the network, all neurons in P1 also received electrical stimulation (black) concurrent with task input (Figure 1). At *t* = 3 s, task-related inputs and stimulation ceased. Decision making experiments show a sigmoidal relationship between coherence and accuracy. To capture this relationship, coherences were sampled logarithmically around the empirically determined center of the sigmoid for each stimulation condition. For control stimulation, this was 0%^2,32^. For pulsatile and cathodic galvanic stimulation, the center coherence was estimated as −57.0%. For anodic galvanic stimulation, the center coherence was estimated as +30.0%.

### Biophysical Attractor Model

The biophysical model was based on a well-established decision-making network^2^. The model simulates a two alternative forced choice task with P1 (blue) and P2 (red) encoding task input (strength of moving dot leftward versus rightward motion). A non-selective (NS) population (yellow) and inhibitory interneuron (Int) population (purple) are also included for a winner-take-all network construction (Figure 1A). The network model consisted of *N* neurons (80% pyramidal neurons and 20% inhibitory interneurons), connected with weights:

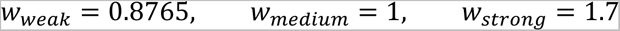

for weak, medium, and strong connections respectively (Figure 1, strength shown with line thickness). Importantly, all neurons in this model were connected with one of these three weights. The model simulated neurons with leaky-integrate-and-fire (LIF) dynamics and synaptic currents from AMPA, NMDA, and GABA receptors. For our simulations, *N* = 1000 neurons and a time step of *dt* = 0.05ms were used.

### Realistic Intracortical Microstimulation

For the sake of computational efficiency, the perceptual decision-making model extended here^2^ approximates individual neural behavior with leaky-integrate-and-fire (LIF) dynamics. Although this simplification yields accurate population-averaged neural firing rates under physiological conditions, it does not naturally accommodate intracortical electrical microstimulation. Therefore, to understand the effects of electrical stimulation on decision making, we incorporated realistic intracortical microstimulation simulation into the Wang model^2^, using a series of approximations to reduce the computational needs of the model. We validated our approach by comparing the output of our modified LIF model to the output of the cable equation model with Hodgkin-Huxley channel dynamics, which is the standard model for assessing extracellular current stimulation on simple neuron morphologies^39^.

The first challenge that arises when representing extracellular current stimulation with LIF neurons is the fact that biological neurons are spatially extended with a gradient of different membrane potentials depending on the location relative to the electrode, whereas LIF neurons only contain a single membrane potential representing the whole neuron. To address this challenge, we note that the magnitude of the voltage response to extracellular current stimulation is greatest in the axon segment that is closest to the electrode (Supplementary Figure 1A). So, we let our LIF neurons approximate only this axon segment, implicitly assuming that stimulation-induced action potentials will be initiated at the segment with greatest voltage response. For extracellular current *pulses*, we approximate the time course of the voltage response as an instantaneous voltage step since pulses induce rapid changes in cable neuron membrane potential (*τ* = 25*μ*s). For *galvanic* stimulation, we inject current such that the voltage response matches cable equation prediction over seconds of stimulation. This approach reliably approximated extracellular current stimulation-induced *voltage* responses in LIF neurons (Supplementary Figure 1B). A final challenge that occurs when using LIF neurons to represent neural spiking activity modulated by electrical stimulation is that spiking responses diverge from voltage responses. To account for this, we varied pulse amplitudes and measured the threshold distance required to elicit an action potential from both cable equation models and our modified LIF neurons. Without any correction, we found that cable equation model thresholds were larger due to the limited duration of the pulse, inhibitory anodic phase, and the inhibitory influence of the side lobes, for which the LIF model does not account. To incorporate these effects into the LIF model, we scaled the magnitude of the LIF pulses with a linear correction factor (k=0.21). After scaling, LIF *spiking* thresholds closely matched those of the cable equation (Supplementary Figure 1C). Finally, we placed 50% of the P1 neurons uniformly around the electrode from 10*μ*m-2mm away based on staining work by Levitt et al.^31^ (1993), and selected pulse parameters to match Hanks et al.^32^: 10*μ*A, 300*μ*s/phase, 200pulse/s.

**Supplementary Figure 1.**
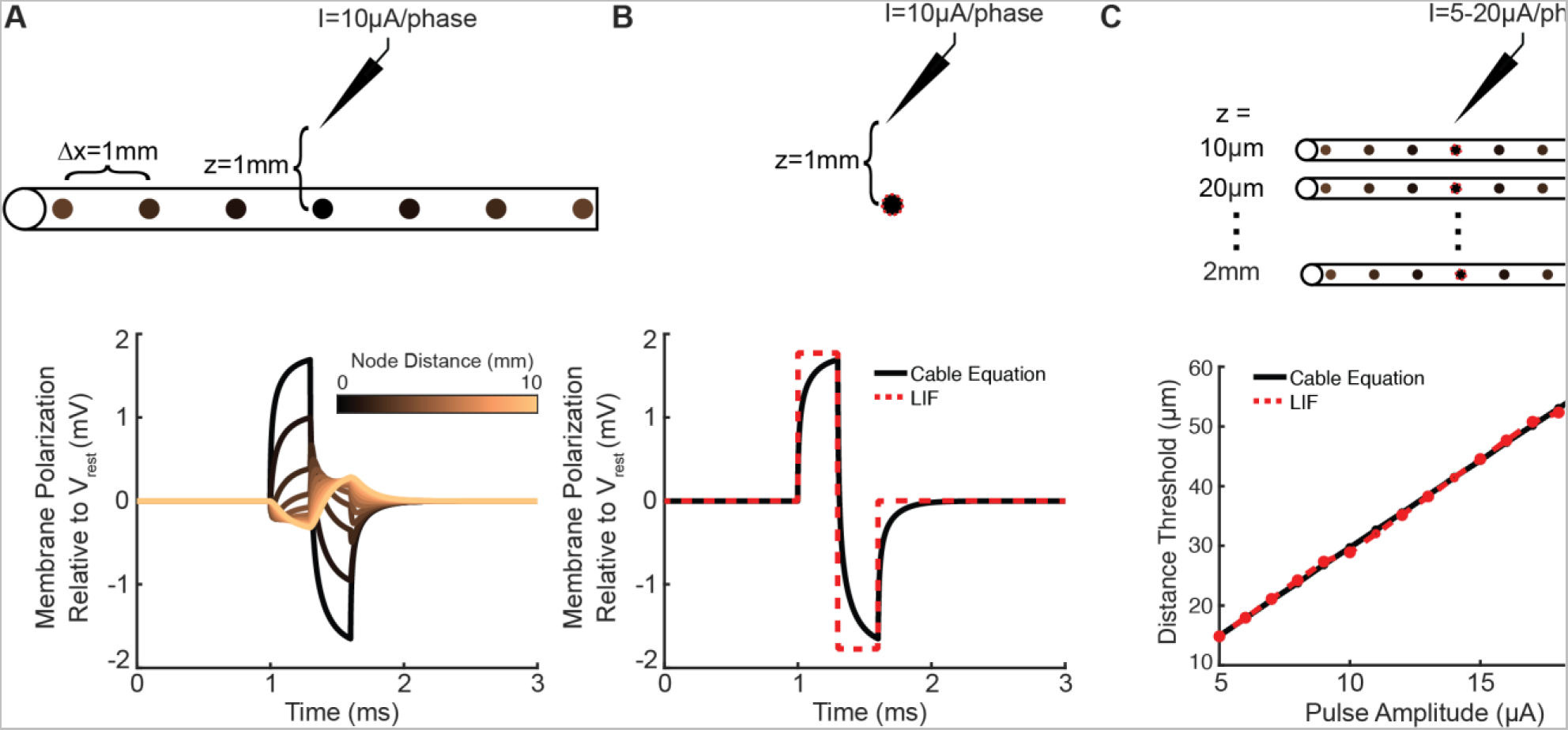
External Current Stimulation in Leaky-Integrate-and-Fire (LIF) Neurons. Cable Equation axon models were subject to biphasic pulsatile stimulation at various distances and amplitudes. (A) A representative voltage response from nodes up to 10mm away from center (axially) is shown for a 10*μ*A/phase biphasic current pulse from a point source located 1mm away (radially). (B) Representative voltage responses from the center node of the Cable Equation model (black) and the LIF approximation (red) are shown. (C) Radial distance required to elicit an action potential from Cable Equation neurons (black) and LIF neurons (red) is plotted for various pulse amplitudes.

### Pulsatile and Galvanic Blocking Effects

In further examination of the previously observed neural responses to pulsatile stimulation^18^, our recent work^19^, using an adapted axon model from Hight & Kalluri^40^, systematically catalogues the possible firing rate responses to biphasic pulsatile stimulation trains. For example, pulses can block spontaneous APs, spontaneous APs can block subsequent pulses, and pulses can block subsequent pulses. These refractory effects are amplitude-dependent, with higher-amplitude pulses causing longer blocking periods, as seen in Supplementary Figure 2A. They are also dependent on the spontaneous firing rate, with higher spontaneous firing rates engendering more pulse-spontaneous interactions, as seen in Supplementary Figure 2C. We therefore expect that delivering PS to a population of neurons would induce a mosaic of activation and deactivation, since each neuron has a different spontaneous rate and experiences a different effective pulse amplitude based on its distance to the electrode. We approximated these effects by adding pulse-pulse and pulse-spontaneous refractory periods like the spontaneous refractory period inherent in the LIF model. To capture the amplitude-dependence, we used blocking times from Steinhardt & Fridman^19^ (0-132ms) recorded at various extracellular amplitudes, using linear interpolation to generate a continuous refractory function.

Using this approach, we replicated the characteristic bends in the pulse rate vs. firing rate curves at various amplitudes (Supplementary Figure 2B) and spontaneous rates (Supplementary Figure 2D). Our approach required temporarily removing the linear correction factor developed in the previous section, likely due to differences in calculating the effects of extracellular current on point neurons between our approach and that of Steinhardt and Fridman.^19^ Also, we observed a substantial decrease in firing rates at high spontaneous firing rates in our LIF-based model (Supplementary Figure 2D) that we did not observe in the adapted Hight & Kalluri model (Supplementary Figure 2C). This discrepancy likely derived from the higher regularity of firing in the LIF model compared to the adapted Hight & Kalluri model. We do not expect this difference to affect our final output since the spontaneous firing rates in the perceptual decision-making network never exceed 50spk/s, above which our LIF approximation is insufficient.

Steinhardt & Fridman^27^ also quantified the depolarizing block that can occur with excessive cathodic galvanic stimulation. This effect is also dependent on the spontaneous firing rate, with higher spontaneous rates inducing block at lower cathodic amplitudes. Similarly, this process yields characteristic bends in the amplitude vs. firing rate curve for GS at various spontaneous rates (Supplementary Figure 2E). We adapted this effect to the LIF model by setting an instantaneous maximum current during GS (1135pA) such that steady state membrane potential could not exceed −19mV based on Qian et al^41^. Anytime instantaneous current inputs exceeded this maximum, the input currents were manually set to 0pA. With this simple rule, we replicated the characteristic bends in the amplitude vs. firing rate curve for GS at various spontaneous rates (Supplementary Figure 2F). Notably, our cathodic block occurred at substantially larger amplitudes than observed in the vestibular model. As with the linear correction factor, this discrepancy likely arises from differences in calculating the effects of extracellular current on point neurons between our approach and that of Steinhardt and Fridman^27^. Importantly, our amplitudes are consistent with those observed in nerve blocking studies^30^.

**Supplementary Figure 2.**
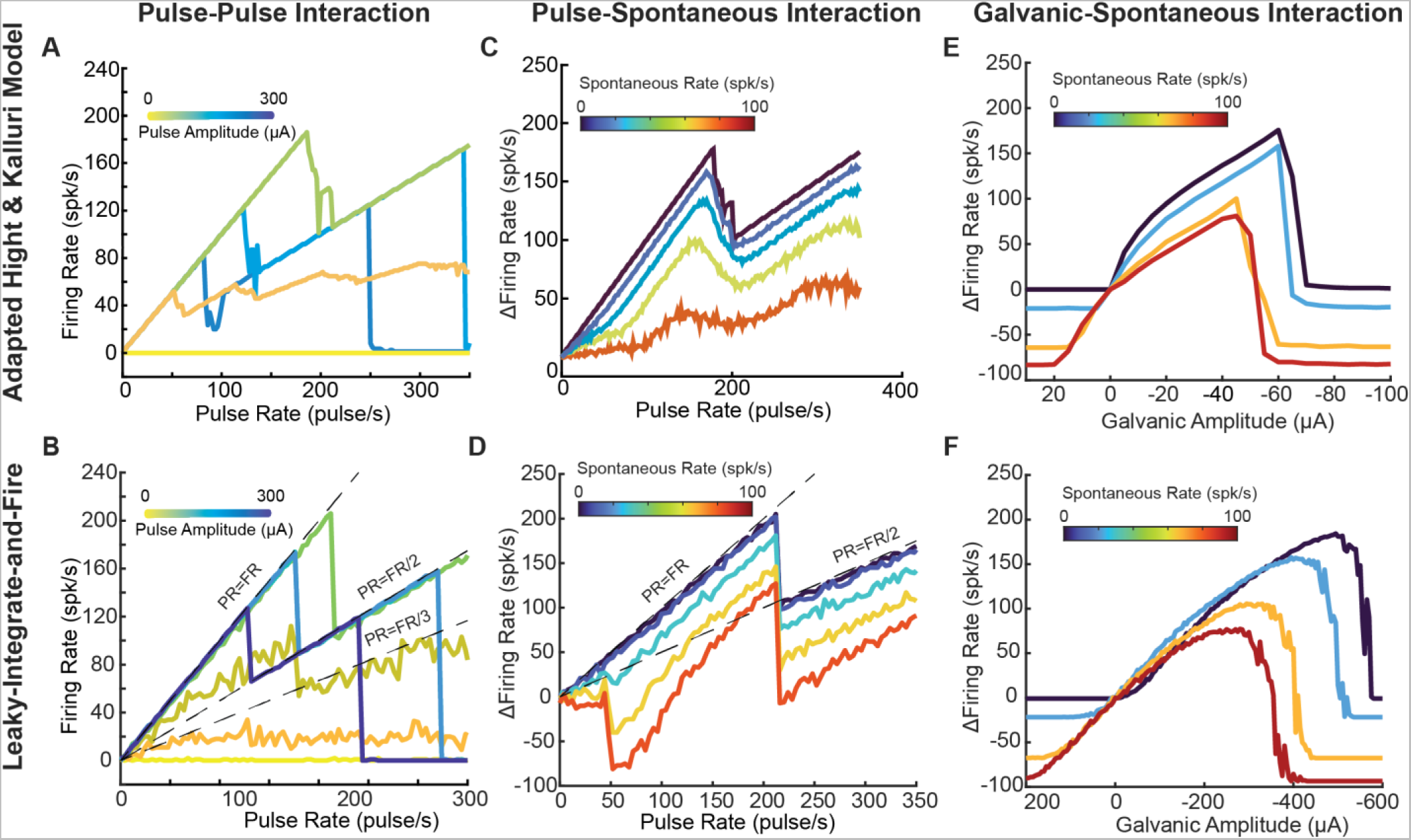
Pulsatile and Galvanic Stimulation Blocking in Leaky-Integrate-and-Fire (LIF) neurons. Neural firing rate in response to various pulse rates and amplitudes are shown for adapted Hight & Kalluri (A) and LIF (B) neurons. Change in neural firing rate from baseline in response to various pulse rates and spontaneous rates are shown for adapted Hight & Kalluri (C) and LIF (D) neurons. Change in neural firing rate from baseline in response to various galvanic stimulation amplitudes and spontaneous rates are shown for adapted Hight & Kalluri (E) and LIF neurons (F).

### Decision Data Analyses

Instantaneous neuron firing rates were calculated in 5ms bins, followed by a 50ms moving average. Population firing rates were then taken as the average instantaneous firing rate of all the neurons in each subpopulation. Decisions were recorded at the end of the 4-second trial, if the final average firing rate of one of the two neural subpopulations (P1 or P2) exceeded 15spk/s, while the other did not. In such cases, the subpopulation whose firing rate exceeded 15spk/s was deemed the “winner” of the trial. This threshold was chosen based on Wang et al^2^. Decision data were then analyzed using logistic regression as in Hanks et al^32^. Differences in decision making were assessed for significance by 2-sample non-parametric bootstrap (N=10,000) on the bias and sensitivity parameters. The time at which the winning subpopulation exceeded 15spk/s after the start of task stimulation (*t* = 1 *s*) was considered the decision time. Trials that reached their decision after the task period (t>3s) were excluded from analyses of time-dependent effects (Figures 4-6). Differences in decision times were assessed for significance by unpaired t-tests.

### Disconnected, Feedback only, and Recurrent Only Network Analyses

Task firing rates were computed by binning all spikes for each P1 neuron between t=1 and t=3 seconds. In the disconnected condition, P1 neurons were not connected to any other neurons in the network. In the feedback only condition, P1 neurons were connected with 30 inhibitory interneurons (keeping an 80/20 E/I ratio) with standard synaptic strength (*w* = 1). In the recurrent excitation only condition, P1 neurons were connected to one another with weakened synaptic strength (*w_rec_* = 0.05). Population-averaged task firing rates were computed by averaging all the individual neuron task firing rates in P1. All stimulation conditions were compared relative to control by subtracting the mean of the control group. Differences in distributions of firing rates across stimulation conditions were assessed for significance by Kolmogorov-Smirnov tests. Differences in individual neural firing rates were assessed for significance by 1-tailed t-tests with Bonferroni correction for multiple comparisons. Differences in population-averaged task firing rates were assessed for significance by Kruskal-Wallis tests.

### Firing Rate Trajectory Data Analyses

Start-of-task firing rates were computed by binning all spikes from P1 neurons occurring between t=1 and t=1.1 seconds and dividing by the duration (0.1s). End-of-task firing rates were computed similarly for the period between t=2.9 and t=3.0 seconds. The firing rate slope around the decision-making threshold (15spk/s) was estimated separately for each trial as

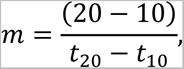

where *t*_10_ is the first time when the P1 population firing rate exceeded 10spk/s and *t*_20_ is the first time the P1 population firing rate exceeded 20spk/s. This method mitigated noise in the P1 population firing rates. The maximum P1 firing rates were computed by finding the maximum instantaneous firing rate in the P1 population for each trial. All stimulation conditions were compared relative to control by subtracting the mean of the control group. All comparisons of P1 firing rate trajectories were assessed for significance by Kruskal-Wallis tests.

### Firing Rate Distribution Data Analyses

Instantaneous neuron firing rates were calculated in 50ms bins, followed by a 200ms moving average. Firing rates were down sampled to 20Hz to match temporal resolution. Kurtosis of the resulting firing rate trace was calculated for each time point and trial separately. Beginning (0-0.5s) and end (3.5-4s) of trials are excluded due to artifactual effects when population firing rates are too low (<1spk/s). Beginning-of-task (t=1.1-1.2s) and end-of-task (t=2.9=3.0s) kurtosis values were obtained by averaging all time points during those time periods. Comparisons of P1 kurtosis were assessed for significance by Kruskal-Wallis tests.

### Spike Timing Data Analyses

Phase-locking of neurons to the pulse stimuli was assessed by measuring the percentage of APs occurring during pulse presentations for each neuron. Regularity of spiking was assessed using coefficient of variation (CV). Synchrony of firing between pairs of neurons was quantified by measuring the percentage of coincident APs, normalized to the neuron with the higher overall firing rate. To facilitate comparison with phase-locking, APs were deemed coincident if they occurred within one pulse phase (300*μs*) of each other. Differences in spike timing were assessed for significance by 1-way ANOVA.

## Supplementary Figures

**Supplementary Figure 3.**
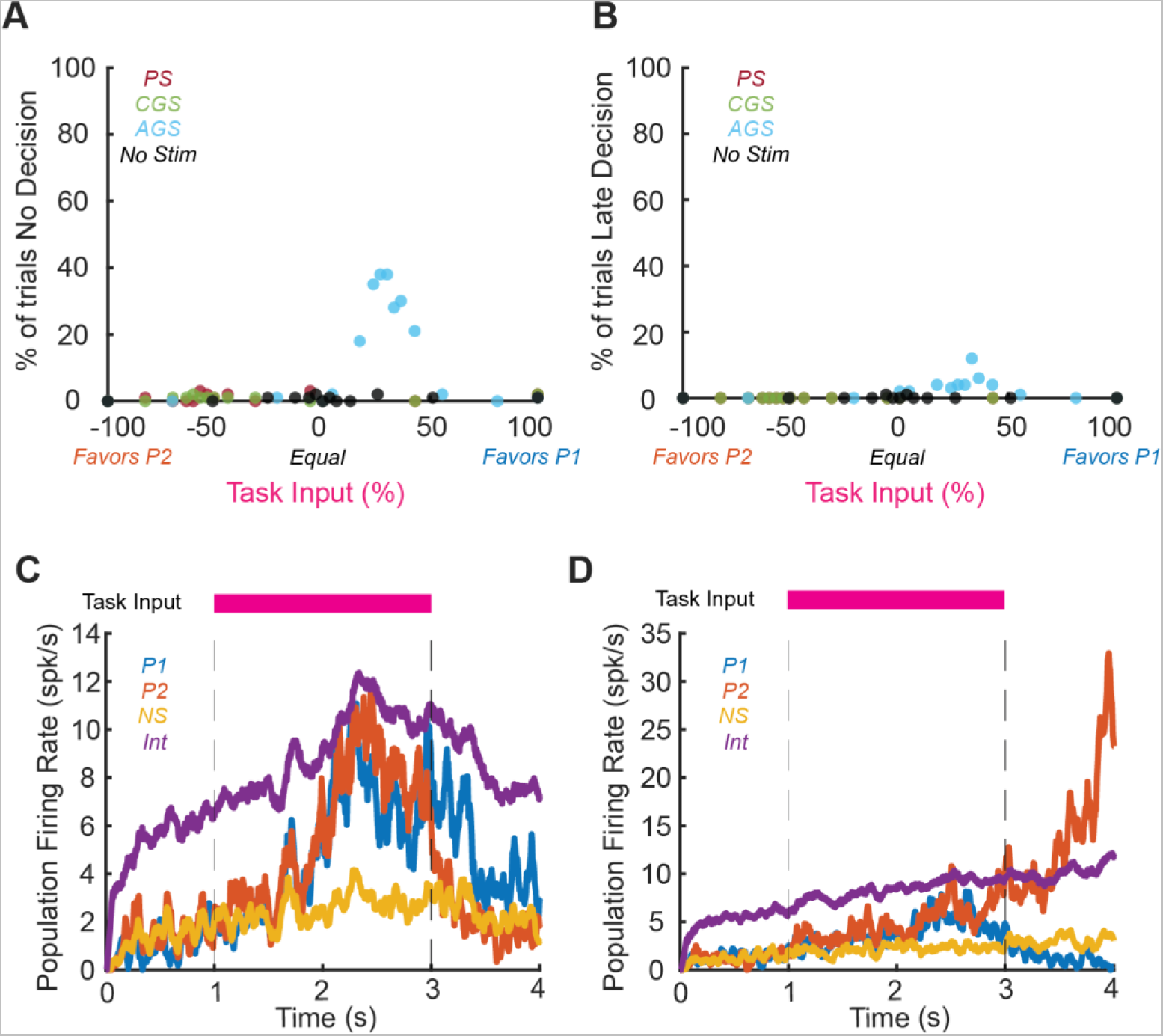
Summary of omitted trials among stimulation conditions (Pulsatile, red; Cathodic Galvanic, green; Anodic Galvanic, blue; and No Stimulation, black) for N=100 trials. (A) Percentage of trials that did not reach a decision is shown as a function of task input. These trials were excluded for all analyses. (B) Percentage of trials that reached their decision after the task period (t>3s) is shown as a function of task input. These trials were excluded from all analyses of time-dependent effects (Figures 4-6). (C) Mean population firing rates of P1 (blue), P2 (red), NS (yellow), and Int (purple) in a representative control trial without electrical stimulation in which no decision was made. (D) Mean population firing rates of P1 (blue), P2 (red), NS (yellow), and Int (purple) in a representative control trial without electrical stimulation in which the decision occurred after the task period.

